# A brain dynamic model based on graph neural network reflect the inter-region interaction of cortical areas

**DOI:** 10.64898/2026.01.26.701662

**Authors:** Shaoxian Li, Debin Zeng, Xiaoxi Dong, Yirong He, Tongtong Che, Jichang Zhang, Zekun Yang, Jie Jiang, Lei Chu, Ying Han, Shuyu Li

## Abstract

A central objective in neuroscience is to elucidate how the brain generates complex dynamic activity through the interactions of brain areas. In this study, we utilized Interaction Network, a graph neural network model, to develop a computational framework for predicting whole-brain cortical blood oxygenation level dependent (BOLD) signals. We derived an Inter-Regional Interaction (IRI) metric to quantify information exchange among brain areas probing the underlying dynamical mechanisms. In addition, the total IRI emitted from each brain region was calculated and defined as the IRI sent by region (RS-IRI). Our model predicted the following 10 time points BOLD activity from initial BOLD signals, and achieved a mean absolute error of 0.04. The predicted functional connectivity (FC) achieves a correlation coefficient of 0.97 compared to the empirical FC. The fluctuation amplitude of the IRI increases with the length of the connection and the largest RS-IRI oscillation amplitude is observed in visual areas. The RS-IRI demonstrates a hierarchical organization, characterized by more concentrated distributions in association regions and larger fluctuation amplitudes in unimodal regions. Applying our approach to Alzheimer’s disease (AD), we demonstrate that the frequency-specific amplitudes of IRI oscillations discriminate AD patients from healthy controls and correlate with Mini-Mental State Examination scores. Together, this work presents a deep learning–based framework for modeling brain dynamics as well a quantitative index of inter-areal interactions, and offers a new perspective for disease characterization.

**Author Summary:** The human brain comprises distinct regions that interact through complex fiber tracts, forming the functional dynamics for diverse cognitive processes. We employed fMRI to assess functional activity and DTI to reconstruct fiber tract connectivity. To elucidate how brain function emerges from these inter-regional interactions, we developed a novel computational framework based on Graph Neural Network (GNN) to model the brain’s interactive dynamics for its capacity to uncover hidden and intricate patterns within data. From this model, we derived a quantitative metric termed Inter-Regional Interaction (IRI), which characterized the fine-grained, dynamic fluctuations in communication between brain areas. Our results suggest that this GNN-based model can accurately simulate brain functional activity and provide a quantitative description of neural interaction patterns. Applying this model to a cohort of Alzheimer’s disease patients, we demonstrated that the IRI metric not only effectively distinguished patients from healthy controls but also significantly correlated with clinical cognitive performance (MMSE scores). This approach advances our understanding of the fundamental principles of brain function and offers a promising tool for identifying the underlying mechanisms of neurological disorders.

## Introduction

The human brain is a highly complex system comprising various types of neurons and glial cells. A central focus of brain research is understanding how the brain generates its intricate dynamic activities, which support a wide range of cognitive functions(1–3). Alterations in these dynamics can indicate systemic brain disorders(4) or transient pharmacological states(5). Many researchers have sought to describe brain dynamics from multiple perspectives, yielding insights into phenomena such as multistability(6–8), metastability(9), co-occurring ripple oscillations(10), and criticality(11,12). A widely accepted consensus is that brain dynamics are markedly nonlinear(13), which are presented at multiple levels, from individual neuron activity to the collective interactions of neuronal populations. This pervasive nonlinearity contributes to the overall complexity of the brain as a system. While these studies analyze and characterize dynamic activities from various perspectives, a pressing question remains: what factors, at both microscopic and macroscopic scales, give rise to these dynamic patterns? A broad proposition suggests that such nonlinear dynamics originate from both local intrinsic activity and inter-regional interactions(14). Crucially, these inter-regional interactions are constrained by the brain’s structural connectivity, which facilitates information transmission between areas and integrates their activities into a globally cohesive yet richly interactive dynamic system, rather than an ensemble of isolated regional dynamics. To investigate the mechanisms underlying these dynamics more sophisticated, mechanism-oriented approaches are needed to explore how these dynamics originate and are shaped.

Dynamical models represent a vital approach for investigating brain dynamics, as their ability to encapsulate complex activities within a single framework enables both the elucidation of underlying mechanisms and the prediction of future states. Since the introduction of the foundational Hodgkin-Huxley (H-H) model(15) and the Wilson-Cowan (W-C) model(16), extensive research has expanded this neuronal framework to encompass larger systems(17). While these studies have successfully constructed dynamic models for individual neurons or specific brain regions, a comprehensive understanding of brain activity mechanisms necessitates the development of dynamic models for the entire brain. Whole-brain dynamical models are typically constructed using network-based modeling, where brain regions are represented as nodes and structural connections between these regions serve as edges. For instance, dynamic causal modeling employs multivariate nonlinear equations to fit regional activity(18–21) and network-based modeling approaches grounded in neural mass models or neural field models have been widely utilized(17,22–25). Several phenomenological models have also yielded significant results in whole-brain modeling. For instance, the Hopf whole-brain model integrates structural connectivity (in the form of adjacency matrices) with local dynamics such as noise and time delays in each brain region (26). This integration enables the analysis and prediction of functional connectivity (FC) (26,27), providing deeper insights into the relationship between brain structure and function(28). Furthermore, they facilitate the simulation of whole-brain activity, allowing researchers to understand various brain states and predict how different regions may respond to external disturbances, serve as valuable tools in pathology and pharmacology research, aiding in the exploration of how brain dynamics react to disruptions or interventions(27). However, these approaches rely heavily on a priori expert knowledge and intuition. We aim to employ a data-driven framework that enables researchers to extract richer insights directly from the intrinsic structure of the data.

In recent years, deep learning has garnered significant attention in neuroscience due to its success in capturing hidden and complex patterns within data. Various convolutional neural networks and recurrent neural networks have proven effective in modeling structural or functional signals (29,30). Motivated by the recognition that the brain can naturally be represented as a graph, there has been growing interest in applying graph neural networks (GNN)(31–33). GNN are specifically designed to treat the brain as a directed graph, where nodes represent anatomical brain regions and edges denote morphological, functional, or structural connectivity between pairwise nodes(34). Regarding node-edge processing paradigms, GNN can be categorized into three primary classes(35): (1) those that aggregate features of neighboring nodes using a learnable filter, as seen in graph convolutional networks; (2) those based on a self-attention algorithm that identifies the most significant neighbors for aggregation, as in graph attention networks; and (3) those employing a message-passing mechanism, where features of both the node are in consideration and its neighboring nodes are combined to learn the local graph representation. In recent years, some approaches have also employed ordinary differential equations as update rules(36,37). Among these approaches, message-passing-based interactive network (IN) enable the prediction of not only the dynamic signal but also the inter-node interactions. In physics, IN have been successfully applied to simulate diverse phenomena(38) and model Newton’s Second Law(39), demonstrating their effectiveness in modeling interaction patterns within complex networks while generating interpretable representations of node interactions.

To achieve a more accurate fitting of whole-brain neural dynamics, this study developed a computational model based on IN to predict BOLD signals across the entire cortical surface. The model calculated interaction information between different brain regions constrained by structural connectivity and predicted dynamic changes in regions through their inter-regional interactions. We trained the model and evaluated its performance on a test set by calculating the Mean Absolute Error (MAE) of the predictions and their correlation with actual FC as the metric of the method’s effectiveness in simulating dynamic variations of cortical signals. We analyzed prediction errors across different regions and investigated its relationship with neural volatility and molecular biological maps to uncover the underlying causes of the spatial distribution in prediction accuracy. Leveraging the model, we extracted inter-regional interaction information at the individual level. By applying this approach to fMRI from Alzheimer’s disease (AD) patients, we found that the model successfully captured oscillatory patterns in inter-regional interactions that differentiate AD patients from normal controls (NC). The constructed model offers a novel approach for analyzing brain dynamics, and the derived interaction metrics provide valuable insights for disease research.

## Results

### Workflow

This study constructed an IN model based on a message-passing mechanism. By fitting and training on BOLD signals, the model generated interaction messages between different brain regions, enabling the simulation of whole-brain dynamics. The model was trained and validated using the resting state fMRI (rs-fMRI) from the Human Connectome Project (HCP) dataset, and the inter-region interactions (IRI) produced by the model were subsequently analyzed. The cortex was segmented into regions according to the MMP360 atlas(40). The mean BOLD signal of each region was calculated as a proxy for the dynamic neural activity within that region (Figure 1A). The specific training strategy is detailed in the Methods section. The trained model was then applied to the test set, and prediction errors were analyzed across different regions (Figure 1B). These error distributions underwent correlation analyses with other brain maps to investigate the sources of regional heterogeneity in prediction accuracy. Subsequently, dynamic interaction messages between regions were extracted, and the information fluctuation maps emitted by different regions were computed (Figure 1C). Finally, we applied our method to data collected from AD patients and control subjects at Xuanwu Hospital (Figure 1D). We compared IRI between the AD and control groups and examined the association between the oscillation patterns of inter-regional interactions and Mini-Mental State Examination (MMSE) scores.

**Figure 1.**
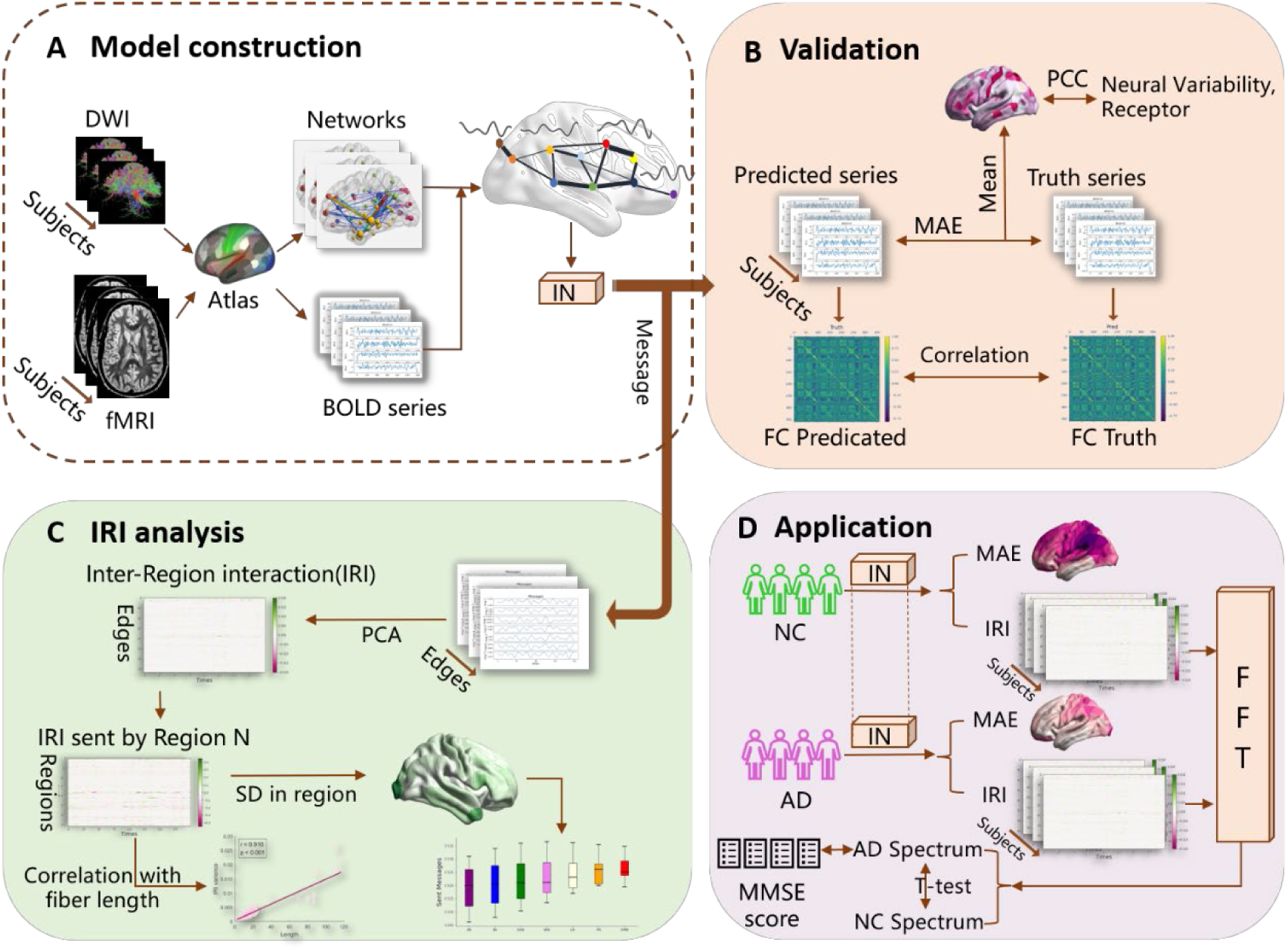
Overall Workflow. **A)** The IN construction based on neuroimaging data and inter-region interaction (IRI) generation. The model was trained independently for each subject, inputting 10 TRs of BOLD signals, and outputting the predicted 10 TRs of BOLD signals. During prediction, the model can calculate the information transmitted between regions. **B)** Validation of the predicted fMRI signals, including MAE and FC correlation analysis. The generated MAE map was further correlated with neural variability and receptor map. **C)** Analysis of IRIs. The message exchanged between each pair of regions, as computed by the model, was reduced via PCA, with the first principal component representing the IRI. The analysis included the IRI on each edge and the total IRI sent by region (see Methods). **D)** Application of the model to AD patients. The IN model for each individual was computed within the AD and control groups to obtain group-average prediction errors and individual IRIs. The IRIs were subjected to fast Fourier transform (FFT) to reveal oscillation patterns in inter-regional interactions. The amplitudes in different frequency bands were then correlated with MMSE scores.

### Model Validation and Prediction Accuracy Analysis

To validate the effectiveness of the methods employed in this study, we constructed individualized cortical neural activity prediction models based on IN for each subject, with data partitioned into training and test sets chronologically, assigning the earlier time points to the training set and the latter time points to the test. Model performance was evaluated using two metrics on the test set: the MAE between predicted and actual signals, and the correlation between FC matrices derived from the predicted and actual signals. The raw BOLD signals were z-scored and the average whole-brain MAE for the predicted data was 0.04. These results indicate that the model can accurately predict dynamic fluctuations across the entire brain. Since each prediction sequence spans 10 TRs, we computed the functional connectivity of the predicted signals using two approaches: a segmented prediction method and a continuous prediction method. The segmented method assesses the model’s ability to reproduce FC over short timescales, while the continuous method evaluates performance in long-term prediction scenarios, where error accumulation may occur (see Methods for details). Both approaches partially recapitulated the structure of the actual functional connectivity (see Figure 2B, Figure S1). The correlation coefficient between the FC derived from segmented predictions and the true FC is 0.9745 ± 0.0067, while the correlation for the continuous prediction method is lower at 0.4957 ± 0.0897.

**Figure 2.**
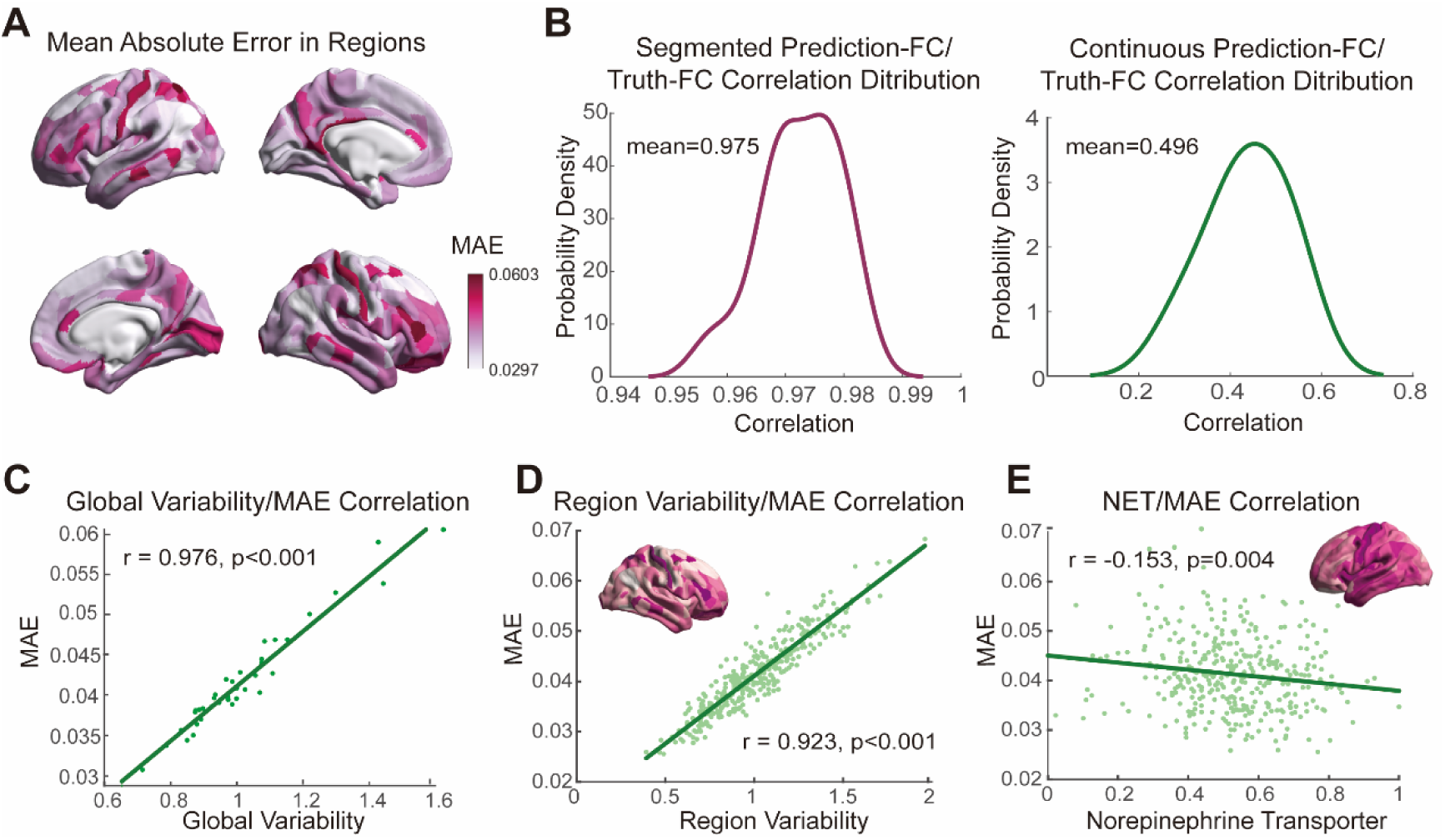
Model validation results. **A)** The distribution across the whole brain of the group-averaged MAE (mean absolute error) for model-predicted fMRI signals in each brain region. Drawn by BrainSpace(105). **B)** Comparisons between FC derived from segmented prediction and continuous prediction with the true FC. The left panel shows the distribution of correlation between the FC obtained from segmented prediction and the true FC. The right panel shows the continuous prediction. The details of continuous and segmented prediction methods are provided in the Methods. **C)** The relationship between MAE and global neural variability. One point denotes a subject. **D)** The relationship between MAE and regional mean neural variability. **E)** The relationship between the distribution of MAE and the distribution of the norepinephrine transporter.

We found that predictive accuracy (MAE) varies across brain regions (Figure 2A). The frontal lobe yields the best predictions and the visual area yields the poorest. We hypothesized that differential prediction performance across brain regions reflects their distinct neural variability. To test this, we examined the relationship between predicted MAE and neural variability. We defined neural variability as the standard deviation (SD) of regional signals(41). Our analysis revealed that, at both the individual and regional levels, the model-predicted MAE was positively correlated with neural variability, supporting the hypothesis that a higher MAE reflects more complex and flexible regional activity (Figure 2C, D), therefore, harder to predict.

To further investigate the underlying factors affecting prediction accuracy across brain regions, we correlated the MAE distribution with a diverse set of cortical maps. We hypothesized that regional activity could be influenced by both the region’s microstructure and mesoscopic functional properties. Therefore, we conducted correlation analyses between the region-wise average MAE maps derived from our model and various maps from neuromaps(42), including functional gradients, coarse-grained structural features, microstructural receptor distributions, and finer-grained gene gradients. All correlations were assessed using permutation tests and corrected for multiple comparisons using the false discovery rate (FDR). Notably, we found a statistically significant but modest negative association between model-predicted regional MAE and the mean distribution of norepinephrine transporters (NET) (Figure 2E), as observed with Methylreboxetine targets(43). These receptors modulate presynaptic sodium uptake, thereby terminating neural signaling(44). This finding may suggest that regions with a higher norepinephrine transporters density leads to greater inhibition of noradrenergic signal transmission, which give rise to more stable signals and enabling more accurate predictions by the model.

### Inter-regional interaction information and its characteristics

The advantage of IN model is the ability to capture information transfer between distinct brain regions. In this study, we derived the IRI from functional signals, utilizing the extraction method described in the Methods section. To address the 30-dimensional interaction messages, we reduced the data by selecting the first principal component as the IRI (Figure S2), resulting in a time-varying IRI sequence for each inter-regional connection (Figure 3A). Also, we aggregated the interaction messages originating from each region and similarly selected the first principal component of this sum as the IRI sent by region (RS-IRI). We used the SD of the IRI as a measure of the flexibility of information transfer between brain regions. A positive correlation was observed between this flexibility and the length of the inter-regional connection (Figure 3B), suggesting that long-range connections may exhibit a greater capacity for adaptive information transfer. This increased capacity may be due to their exposure to more dynamic transmission contexts, the involvement of more complex signals, and the need for larger signal fluctuations to ensure accurate transmission.

**Figure 3.**
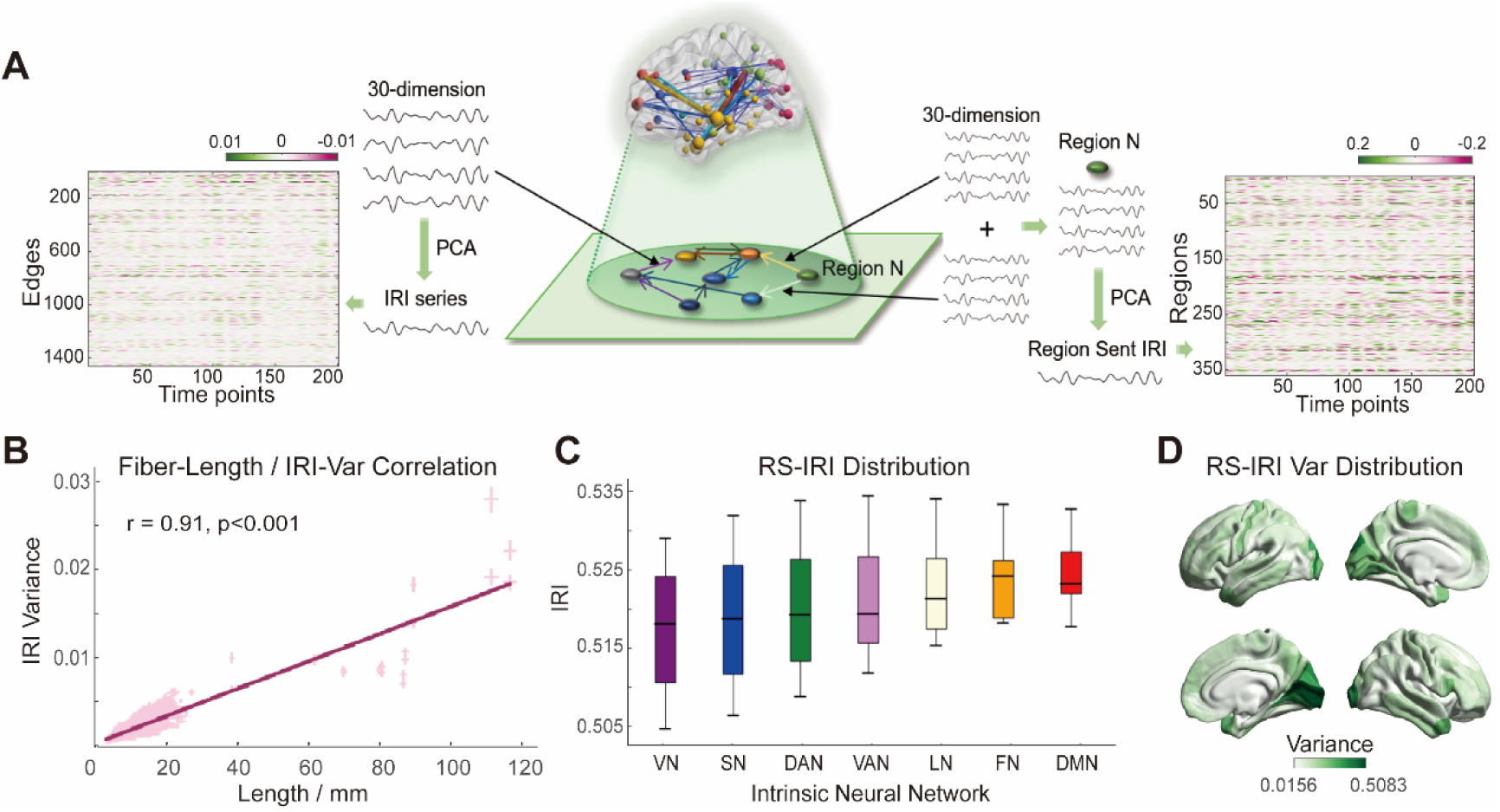
Analysis of inter-regional interaction information (IRI). **A)** Flowchart for IRI derivation: Initially, 30-dimensional time series served as interaction messages for each connection. Dimensionality reduction was performed to derive the IRI time series. IRI was quantified for each fiber tract, and the sum of IRI sent from each brain region was termed IRI sent by region (RS-IRI). **B)** Correlation between IRI variance and fiber length. The error bars for each point represent the standard deviation across subjects for the corresponding connection length and IRI variance. **C)** Distribution of RS-IRI strengths across different subnetworks. **D)** Distribution of RS-IRI variability.

To investigate whether different functional sub-networks send information with distinct strengths, we plotted boxplots of emission strength across sub-networks (Figure 3C). We found that as hierarchical level increases, the distribution of emitted information strength exhibits lower variance. This centralization may arise from higher-order regions, such as the default mode network (DMN), exhibiting a relatively lower excitation–inhibition ratio, which leads to stronger inhibitory signals and consequently more stable information emission(45,46). Finally, we quantified the variance of RS-IRI across brain regions, finding that visual regions exhibited the greatest flexibility in emitting interaction information (Figure 3D). This may reflect the highly diverse information generated and output by these regions.

### Application in Alzheimer’s diseases

To investigate whether neurodegenerative pathology induces alterations in the dynamic interactions among cortical regions, we applied our computational model to data from AD patients. Our analyses revealed differences in the mean predicted MAE between the AD and NC groups but did not show significant group differences (Figure 4A), likely due to the limited sample size. Before computing the IRI, we compared the SD of interaction messages across each dimension between the AD and NC groups, which we used as a proxy for the amount of information carried by the IRI. We found that, in most dimensions, the between-group differences were significant, while non-significant dimensions tended to have lower SD, suggesting that these dimensions may carry less informational content (Figure 4B). The SD of interaction messages in the NC group was significantly higher than in the AD group, indicating greater flexibility in inter-regional interactions for the NC group. This may reflect that the AD patients experience dysfunction in inter-regional information transfer with less informative content.

**Figure 4.**
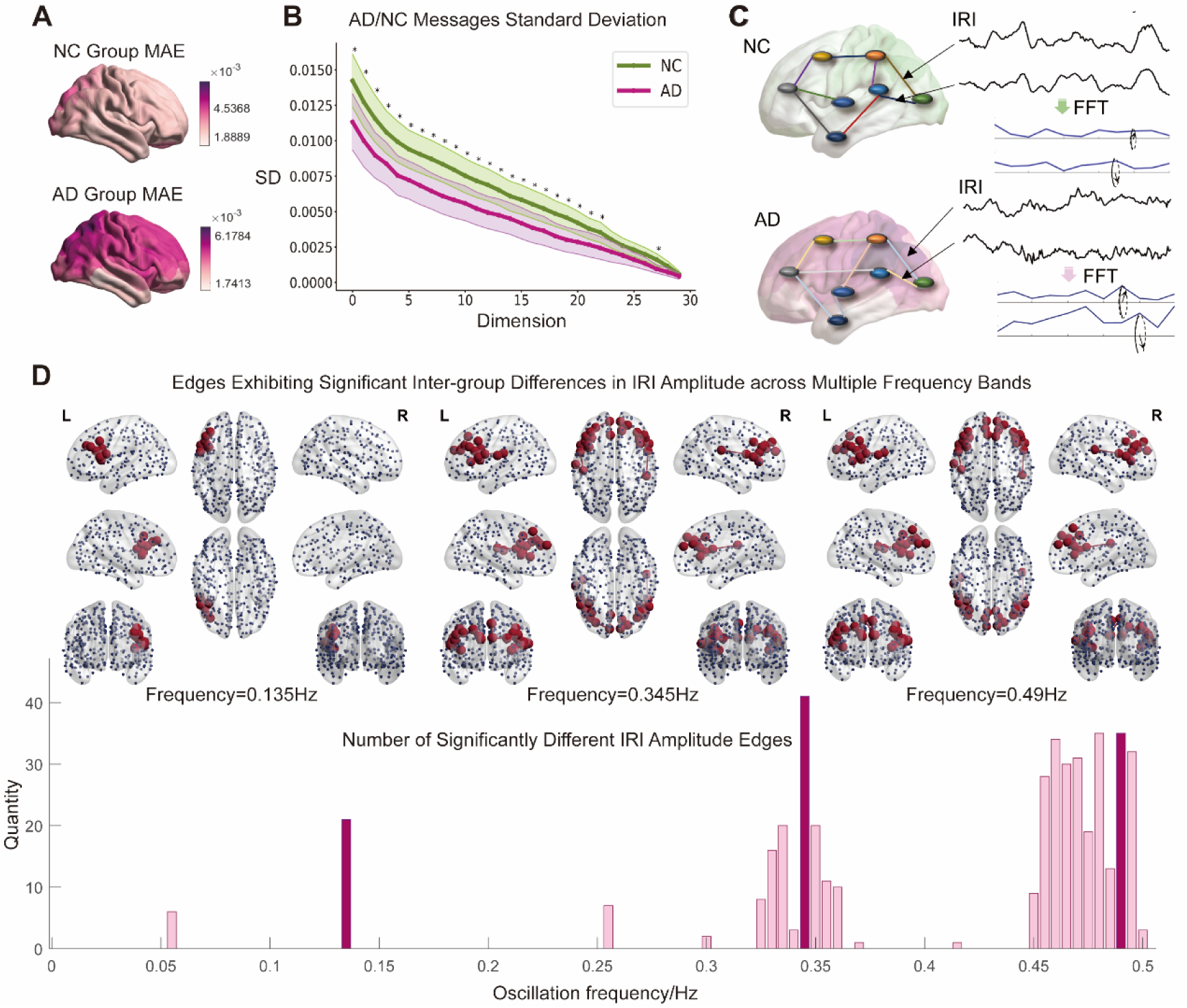
Group-level comparison between AD patients and controls. **A)** The group-averaged prediction MAE maps on the cortical surface for the AD and NC groups. **B)** The between-group differences in the SD across each dimension of the interaction messages for AD and NC groups, with asterisks indicating statistically significant between-group differences at α = 0.05. **C)** The brief workflow for obtaining oscillation amplitudes across different frequency bands of the IRI in the AD and NC groups. **D)** The between-group differences in the oscillation amplitudes described in (A). For each frequency band, the IRI between-group differences are computed and subjected to FDR correction. The top figure shows, at low-, mid-, and high-frequency ranges (0.135, 0.345 and 0.49Hz), the connections exhibiting between-group differences in IRI oscillations and the corresponding brain regions, drawn by BrainNet Viewer(106). The chosen frequency bands correspond to the deep purplish-red bands shown in the figure below. The bottom figure presents the total number of connections between brain regions that show significant between-group differences in oscillation amplitudes across different frequency bands for the IRI.

We computed the IRI for each subject in both the AD and NC groups. Our results indicated that the AD and NC groups differ in their IRIs, with the AD group showing more high-frequency components (Figure 4C). To examine whether the spectral content of the IRI could distinguish AD patients from controls, we performed Fast Fourier Transform (FFT) on the IRI signals for each edge, obtaining their spectral profiles. We then conducted t-tests on the amplitude at each frequency band, applying FDR correction, the results see supplementary table. Results confirmed our hypothesis, revealing significant differences in IRI predominantly in the high-frequency range, where the AD group exhibited markedly increased IRI on connecting edges (Figure 4D). This attenuation of high-frequency inter-regional information exchange suggests that cognitive deficits in AD may be associated with increased high-frequency noise contamination in neural interactions. We analyzed the connections showing significant differences in IRI across different frequency bands and found that the majority of IRI with significant differences were located in the information flows of intra- or inter-higher-order regions, particularly involving the frontoparietal network and the ventral attention network (Figure 4D).

We further investigated the relationship between IRI across different frequency bands and cognitive impairment in AD patients, we correlated the amplitude values of each edge within each frequency band with the MMSE scores in the AD group. Significance was evaluated using permutation testing with FDR correction. Ultimately, we identified six edges where amplitude of communication in specific frequency bands significantly correlated with MMSE scores, predominantly distributed in the medial and lateral prefrontal cortices and the cingulate cortex (Figure 5). All these regions are situated within the fronto-parietal network, default mode network, and ventral attention network, indicating that IRI within and between these networks may reflect cognitive performance as measured by the MMSE in AD patients.

**Figure 5.**
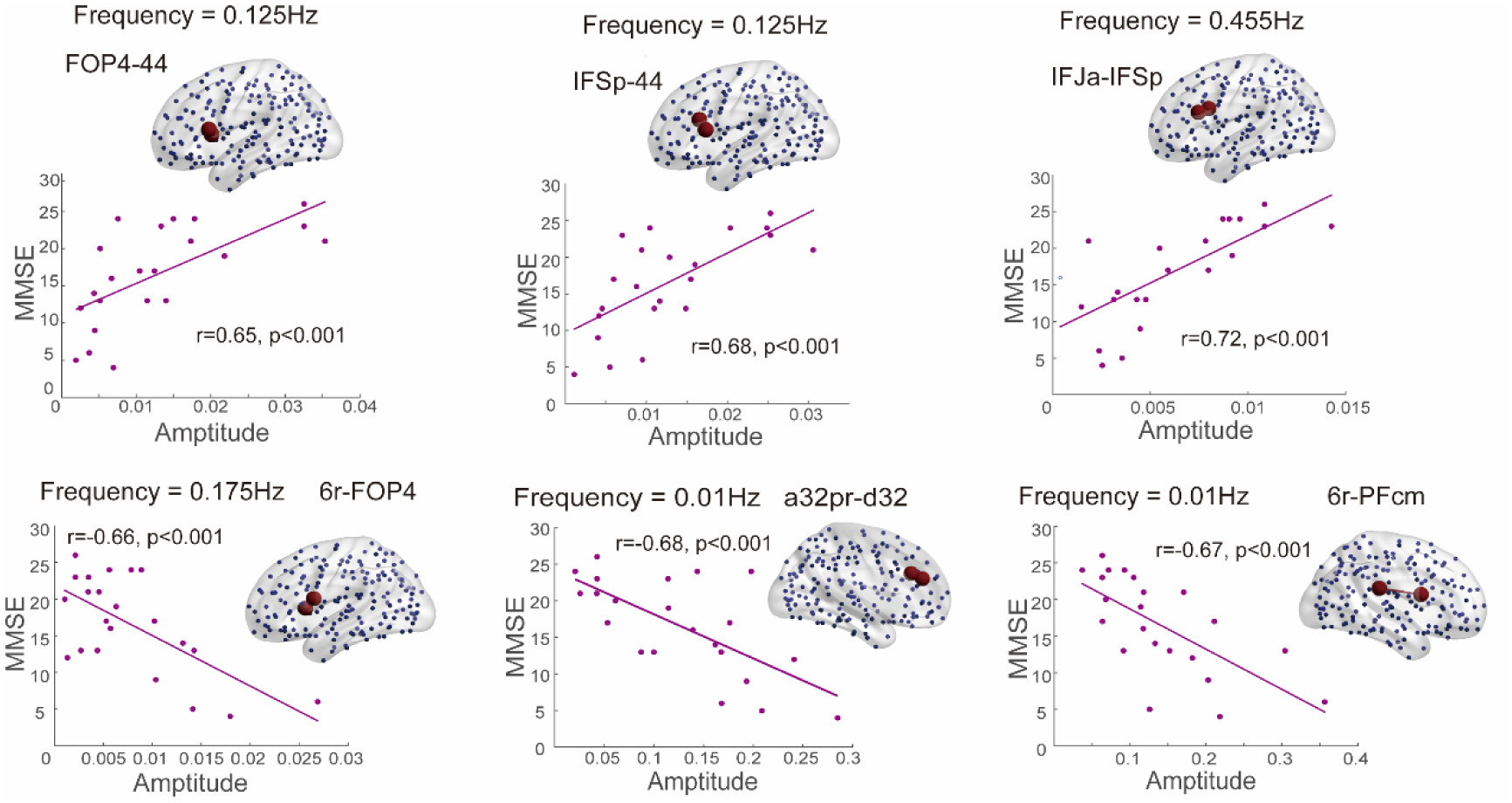
The association between oscillation amplitude across different IRI frequency bands and MMSE scores in AD patients. In this figure, the connections and their connecting brain regions for which IRI is significantly correlated with MMSE scores. A permutation test was used, with all results FDR-corrected. The plotted fit line represents a linear (first-order) fit. The connections corresponding to the IRI oscillation amplitudes are projected onto the brain, with the associated connections annotated at the top or on the right side. The numerical labels adjacent to the brain correspond to the indices defined in the MMP atlas. Connected regions are separated by a hyphen (e.g., ‘ID-ID’). For a complete list of MMP region IDs, see supplementary.

## Discussion

Our study developed a computational model based on IN to predict BOLD signals across the entire cortical surface. This model computes interregional interaction information constrained by structural connectivity and predicts dynamic BOLD changes in brain regions through their inter-regional interactions. Validation using MAE and FC correlation demonstrated that this approach effectively simulates dynamic variations in cortical signals. By extracting inter-regional interaction information at the individual level from the model and applying it to a dataset of AD patients, we found that the model captures distinct oscillatory interaction patterns between brain regions in AD patients compared to NC. The constructed model offers a novel method for analyzing brain dynamics, and the derived interaction metrics provide new perspectives for disease research.

### The rationale for using GNN in brain dynamical modeling

GNN represent a deep learning framework adept at analyzing network-structured data. They are extensively applied across various domains, including power grids(47), transportation networks(48), and financial systems(49). Recently, some researchers have begun employing GNN for modeling neural systems(50,51) due to their superior capacity for processing and analyzing network-structured data. Most GNN utilize graph convolution modules(52–54), while others use message-passing mechanisms, which, although computationally intensive, can capture interactions between pairs of nodes, such as gravitational interactions among planets(39) and forces between high-energy particles(55). This message-passing approach emphasizes connections but requires greater computational capacity. To enhance the reliability of interaction information and minimize interference from multiple computational modules, this study adopts a streamlined model construction approach: both the interaction information generation module and the node state generation module are constructed using simple multilayer perceptron (MLP), without integrating additional computational components.

Inspired by these studies, we developed an IN model capable of predicting cortical BOLD signals by leveraging dynamic interregional interactions. The model achieved an average MAE of 0.04 in predicting overall fMRI signals. This results represent an improvement over the diffusion convolution recurrent neural network (DCRNN) and graph WaveNet (GWN) used in prior research(53), demonstrating the model’s effectiveness in capturing dynamic fluctuations in cortical BOLD signals.

The segmented prediction approach facilitated partial reproduction of FC structures (see Figure 2C, D), with a correlation coefficient of 0.9745 to actual FC, slightly higher than the 0.95 reported for DCRNN(53). However, due to different computational methodologies, these values are not directly comparable. Consequently, the correlation between sequentially predicted FC and true FC, affected by accumulated errors, was lower at 0.4957. This result, like other data-driven models(30), is modest compared to most traditional dynamical models, which typically begin with empirical equations and structured networks and iteratively tune parameters to enhance the similarity between generated and actual FC(27,56,57). Improving FC correlation in generated sequences is plausible since these sequences need only replicate certain structural aspects of FC rather than its full range of features. Our method optimized the model by fitting it to real sequences, minimizing the discrepancy between generated and actual data via the loss function. Despite the model’s underwhelming performance in continuous FC prediction, which indicates a deficiency in capturing long-range temporal dependencies, it enables derived inter-regional interactions to inform subsequent predictions of cortical dynamics. Ideally, if the model could perfectly simulate whole-brain dynamics, the generated sequences should exhibit a complete FC structure. Unfortunately, the current model appears focused primarily on input and target 10 TR data, aligning with our design objectives, while neglecting long-range influences. This limitation is understandable, as such constraints were not explicitly incorporated into the model. Prior studies have suggested that long-range BOLD signal simulations driven by data depend on noise(30). Since our model excluded noise, long-range predictions tend to converge toward attractors over time. We believe, however, that introducing noise could undermine the reliability of the IRI metrics, leading us to incorporate only regularization-based noise introduction, avoiding explicit noise-driven modeling.

### Understanding and application of IRI

Using our model, we extracted IRI and related it to the concept of information flow. Historically, information flow has been described as the directionality and strength of information transfer among different brain regions(58,59). This definition, in conjunction with effective connectivity, bears similarity to approaches such as dynamic causal modeling(60). Recent advancements in DTI have provided a structural basis for information flow(61,62). However, characterizing information transfer directionality in a purely static manner is insufficient. To fully understand information processing in the brain, including perception, transduction, coordination, storage, and generation, an analysis of dynamic fluctuations is imperative, as these processes reflect continuous changes in the brain’s underlying states, even under constant environmental conditions(63,64). These dynamics evolve over time and are rooted in the brain’s structural connectivity(65). Therefore, when constructing a comprehensive brain communication model, these factors must be duly considered(66,67). In this study, the whole-brain dynamical model based on GNN generates dynamic interregional information transfer by integrating BOLD signals from different regions and their structural connectivity, representing these processes as a temporal sequence. This sequence is akin to edge Time Series (eTS), originally defined as a co-fluctuation time series, the pointwise product of z-scored signals from two brain regions at successive time points(68). It provides a mechanism for deconstructing FC into temporal sequences. A connection-centric measure called edge functional connectivity (edge FC or eFC) captures the dynamic expression of brain functional systems through fine-scale edge time series analysis(69,70). However, unlike eTS, the IRI produced from the IN model is not reconstructed from static FC, thereby avoiding concerns regarding its “dynamic” nature(71–73). Crucially, the IRI is grounded in the underlying structural connectivity derived from DTI, rather than being inferred solely from functional signals. Essentially, IRI offers a method for deriving fluctuations in connection edges directly from dynamic variations in fMRI signals, and these fluctuations are both structure-based and biologically grounded.

We observed a positive correlation between the variability of IRI and the length of connections. Long-range connections are known to play a crucial role in brain activity(74). Although some studies suggest that brain geometry alone can reconstruct functional connectivity(75), other research, both experimental and clinical, demonstrates that sparse long-range connections are vital for the complexity of brain functional dynamics(76–78). Our findings support this notion, indicating that long-range connections are essential in whole-brain modeling, with their significance increasing with connection length. Additionally, some studies report that distant projections are more strongly influenced by electrical stimulation(79), which may explain our results: greater IRI variability could correspond to a higher information-carrying capacity, suggesting that long-range connections transmit more potent electrical stimuli. The average strength of IRI emitted by different brain regions exhibits a gradient distribution across various functional subnetworks, indicating that higher-order brain regions generate the strongest IRI during resting-state conditions. This suggests that these regions are more likely to play a dominant role in resting-state activity.

In the analysis of AD patients, we noticed that the IRI in the AD group contained more high-frequency components, yet the standard deviation (SD) of the interaction messages was significantly lower. This suggests diminished flexibility in information transmission coupled with increased erroneous noise, which reduced the proportion of meaningful content within the transmitted signals. We observed significant differences in the IRI within the 0.4–0.5 Hz frequency band compared to NC. These oscillatory pattern variations have been documented in EEG studies(80). Unlike previous research that focused on signals from individual brain regions, our study employs IRI; since IRI is inherently related to the underlying regional signals, the findings align with earlier research indicating abnormal neuronal firing activity in the theta frequency band in AD patients(81). Due to the limited temporal resolution of fMRI data, we cannot capture information in higher frequency ranges. Nevertheless, leveraging the high spatial resolution of fMRI allows us to analyze how oscillatory interaction frequencies between brain regions relate to cognitive scores and to identify specific connection sites involved. We identified six connections where IRI significantly correlates with MMSE scores in AD patients, predominantly distributed in the medial and lateral prefrontal cortices and the cingulate cortex. Notably, four of these links connect the DMN to other networks. These findings support prior research on functional connectivity alterations in AD patients(82), reinforcing the notion that the DMN exhibits pathological changes in AD, subsequently affecting other regions involved in higher cognitive functions via interregional interactions.

### Limitations

First, this study employs GNN to predict discretized fMRI signals. However, it does not construct continuous dynamic models akin to differential equations. Future research incorporating mechanistic approaches from machine learning(83) could facilitate the development of brain dynamics models capable of continuous prediction. Second, the structural connectivity in this study was based on data obtained through DTI tractography for model construction. However, DTI imaging may have accuracy limitations and often overlooks non-long-range synaptic transmissions between brain regions (e.g., adjacency effects). The brain’s dynamic behavior may not solely depend on network-like structural connectivity(84); some studies suggest that brain dynamics could also arise from geometric morphology(75). Integrating approaches such as geodesic distance(85) and cellular architecture similarity networks(86) can complement these various connections. Employing heterogeneous GNN methods(87) may aid in constructing such multifaceted models. Third, current models are primarily data-driven at a macroscopic scale and neglect microstructural elements such as neurons. They also do not incorporate task-driven models at the phenotypic cognitive level. Future work should aim to integrate task-based approaches, developing multimodal, multiscale models based on microscopic priors, aligning with the goal of multiscale brain modeling(88).

## Methods

### Participants

This study utilizes two batches of data, obtained from the Human Connectome Project (HCP) by the National Institutes of Health (NIH)(89) and from the Neurology Department at Xuanwu Hospital, Capital Medical University in China. Following quality control procedures based on head motion and the length of the remaining time series, a total of 148 participants from the HCP dataset were retained, including 72 males and 76 females, aged between 22 and 35 years. In addition, we enrolled 41 patients diagnosed with Alzheimer’s disease (AD) from the memory clinic of the Neurology Department at Xuanwu Hospital, Capital Medical University (CMU) in China. We also recruited 42 healthy individuals as normal controls from local communities in Beijing, China. This study obtained approval from the Medical Research Ethics Committee at Xuanwu Hospital, and all participants provided written informed consent. Standard clinical assessments were conducted on all participants, which included a thorough medical history investigation, neurological examination, and neuropsychological tests. The cognitive tests administered encompassed the Montreal Cognitive Assessment (MoCA, Beijing version)(90), Auditory Verbal Learning Test (AVLT), Clinical Dementia Rating (CDR)(91), Mini-Mental State Examination (MMSE), Hamilton Depression Rating Scale (HAMD), Activities of Daily Living (ADL) scale, Hachinski Ischemic Scale, and the Center for Epidemiologic Studies Depression scale(92). The patients with AD were diagnosed based on the criteria of the National Institute of Aging-Alzheimer’s Association (NIA-AA) for clinically probable AD(93). After data preprocessing, the final sample comprised 34 healthy control individuals and 23 AD patients. The age range of the participants was between 53 and 81 years. Among the AD group, there were 14 males and 9 females, while the control group consisted of 21 males and 13 females.

### Data acquisition

The HCP dataset comprises resting-state functional magnetic resonance imaging (rs-fMRI) and diffusion magnetic resonance imaging (dMRI) data. Each participant in the HCP dataset underwent two 15-minute resting-state fMRI scans at different time points, acquired from two directions. This resulted in a total of 60 minutes of rs-fMRI data, comprising 4800 repetitions with a Time Repetition (TR) of 720ms. The scans were performed on a Siemens Skyra scanner with a 3T field strength. This study utilizes two batches of data, obtained from the Human Connectome Project (HCP) by the National Institutes of Health (NIH)(89) and from the Neurology Department at Xuanwu Hospital, Capital Medical University in China. The HCP dataset comprises rs-fMRI and dMRI data. Each participant in the HCP dataset underwent two 15-minute rs fMRI scans at different time points, acquired from two directions. This resulted in a total of 60 minutes of rs-fMRI data, comprising 4800 repetitions with a Time Repetition (TR) of 720ms. The scans were performed on a Siemens Skyra scanner with a 3T field strength.

The data from the Neurology Department at Xuanwu Hospital, Capital Medical University (CMU), comprises rs-fMRI and DTIdata. All images were acquired on 3.0 T Siemens system (Magnetom Trio Tim; Erlangen, Germany) at the Department of Radiology, Xuanwu Hospital, Capital Medical University, Beijing, China. T1-weighted images were acquired using a magnetization prepared rapid gradient echo (MPRAGE) sequence (TR = 1900 ms; TE = 2.2 ms; TI = 900 ms; flip angle = 9◦;FOV= 224 mm × 256 mm; matrix size = 448 × 512; number of slices = 176; slice thickness = 1 mm). Functional images were acquired axially using a gradient-echo EPI sequence (TR = 2000 ms; TE = 40 ms; flip angle = 90◦; FOV= 240 mm × 240 mm; matrix size =64 × 64; number of sections = 28; section thickness = 4 mm; voxel size = 3.75 × 3.75 × 4mm3;gap= 1 mm; volume number = 239). DTI images were collected axially by using a single-shot echo-planar sequence (repetition time (msec)/echo time (msec), 11000/98; flip angle, 90◦; field of view, 256 × 232 mm2; 128 × 116 matrix; 60 sections; section thickness, 2 mm; voxel size, 2 × 2 × 2 mm3; 30 gradient directions with b value of 1000 sec/mm2 and one image with a b value of 0 sec/mm2; and three averages).

### Data preprocessing

For the HCP dataset, an affine transformation was initially applied to register individual dMRI images with the T1-weighted image. Then, the structural image of each individual was registered to the ICBM152 template(94), yielding an inverse warping transformation from the standard space to the dMRI space. The HCP MMP 1.0 atlas(40) was then mapped onto the sample space using this inverse transformation. This division divided the entire brain into 360 brain regions, with 180 regions per hemisphere. Fiber tractography was subsequently performed using DSI-Studio(95). By calculating the eigenvalues and eigenvectors of water molecules in three directions, fiber tracts were traced. Two weighted matrices, the connection density matrix and the connection length matrix, were constructed based on the fiber tracts between two nodes.

Regarding fMRI data processing, the HCP-pipeline was initially applied for minimal preprocessing(96). Subsequently, a high-pass filter with FWHM greater than 2000s was applied to the fMRI signals and the ICA-FIX(97,98) method was then employed to remove noise components from the fMRI signals, followed by regression of the data using the global signal as a regressor. Finally, a bandpass filter ranging from 0.009 to 0.08 Hz was applied to eliminate most of the noise. After post-processing, the entire cerebral cortex was segmented according to the HCP MMP 1.0 atlas(40). The average fMRI signal within each brain region was calculated, obtaining fMRI signal values for the 360 brain regions. Incomplete data were discarded, and subjects with head motion exceeding 0.25 were excluded. Among the remaining subjects, time points with head motion exceeding 0.25 were removed, and the data were divided into non-uniform intervals. Each interval overlapped by grouping them in sets of 20 time points, where the first ten time points represented the input data, and the latter ten time points represented the true values of the output data. If an interval contained fewer than 20 time points, it was discarded. The data from individuals with input-output groupings greater than 2000 were retained, resulting in a total of 148 participants, including 72 males and 76 females, with ages ranging from 22 to 35 years.

As for the data from the Neurology Department at Xuanwu Hospital we utilized the MRTrix 3.0 pipeline(99) for DTI preprocessing. Firstly, an affine transformation was applied to register individual DTI images with the T1-weighted image. Subsequently, the structural images of each individual were registered to the ICBM152 template(94). Computing the eigenvalues and eigenvectors of water molecules in the brain along three directions to trace the fiber and obtained connection probabilities. The connection probabilities between two brain regions were summed based on the HCP MMP 1.0 atlas(40), resulting in a connection probability-weighted structural connectivity matrix (SC).

We utilized the fMRIPrep 20.0.7 pipeline(100) for the preprocessing of fMRI data. The preprocessing steps for the T1-weighted images included intensity non-uniformity correction, skull stripping, spatial normalization to the standard space (MNI152NLin6Asym), brain tissue segmentation, and cortical surface reconstruction. The preprocessing steps for the rs-fMRI data involved removing the first five time points, motion correction, slice-timing correction, and co-registration to the corresponding T1-weighted images using boundary-based registration. Subsequently, we employed spatial smoothing using an isotropic Gaussian kernel with a full width at half maximum (FWHM) of 6 mm. In the denoising phase, we performed automatic removal of motion artifacts using ICA-based Automatic Removal Of Motion Artifacts (AROMA)(101), a 24-parameter model, and linear regression, which included signals from the mean white matter and cerebral spinal fluid(102). This was followed by the application of a high-pass filter using a discrete cosine filter with a cutoff period of 128 s. After processing, the whole-brain cortex was parcellated according to the HCP MMP 1.0 atlas(40) and the average fMRI signal within each brain region was calculated to obtain signal values for 360 brain regions. After excluding the images with significant head motion (head motion exceeding 0.5), the final sample comprised 34 healthy control individuals and 23 AD patients. The age range of the participants was between 53 and 81 years. Among the AD group, there were 14 males and 9 females, while the control group consisted of 21 males and 13 females.

### IN models construction

The model can be considered a hidden function *F*(*x*), denoted as *F_alg_*(*x*), and its overall operation process can be represented by an equation.

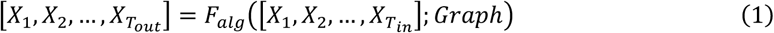

In which *X*_*out*_ represents the predicted fMRI signal, expressed as a 360-dimensional vector, with the subscript number indicating the number of time points. This study uses fMRI signals from 10 time points as input, hence *T* = 10. *Graph* denotes the whole-brain network, which is represented using a structural connectivity matrix derived from dMRI tracking.

The process of information interaction includes two parts: the transmission and reception of information. Based on this idea, *F*(*x*)can be decomposed into two functions: the information interaction function, *I*(*x*), and the state change function, *R*(*x*). This model can be simplified to the level of brain regions. Information is transmitted between brain regions through the interaction function *I*(*x*), as shown in equation (2).

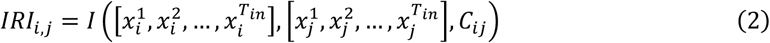

*IRI*_*i*,*j*_ represents the interaction between brain region *i* and brain region *j*, with *C*_*ij*_ being the characteristic of the connection between brain region *i* and brain region *j*, which is estimated with structural connectivity matrices generated from structural connectivity (SC) and global parameter *G*, defined as 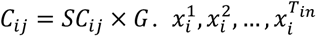 represents the input fMRI signal from region *i* in time *T*_*in*_ = 10. Assuming that the information transfer between brain regions is additive, meaning that for brain region *i*, the influences from different brain regions are directly summated, then the change in state of a brain region can also be expressed in a simplified form through a state change function, equation (3).

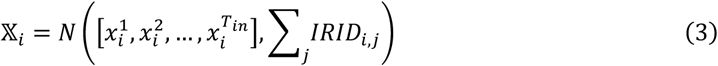

In which 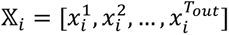 represents the predicted fMRI signal from region *i* in time *T*_*out*_ = 10. To minimize the complexity of the model and enhance its interpretability, this study utilizes the MLP to construct functions *I*(*x*) and *R*(*x*), that is, by using simple linear layers and activation functions for construction. The method of constructing *I*(*x*) involves multiplying the characteristics of the connecting edges directly by the output of the multilayer perceptron, allowing the characteristics of the connecting edges to influence the information interaction between brain regions, which can be represented by the equation (4)

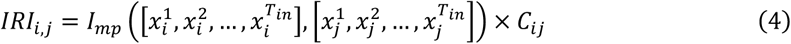

In which *I*_*mp*_ represents the MLP for *I*(*x*).

The model inputs are the values of whole-brain fMRI signals for 10 time points, with the output being the predicted values of whole-brain fMRI signals for the next 10 time points. For each sequence of an individual’s scan, input values are constructed as *input* = 𝕏^*in*,*k*^ *for k* = [0,1,2, …, *k*_*max*_], and truth values as *truth* = 𝕏^*true*,*k*^ *for k* = [10,11,12, …, *k*_*max*_]. 𝕏^*k*^ represente the input fMRI signal in 10 time points, which means a 10×360 dimension vector. Therefore, the number of time points for both input and truth values remain the same, equal to the number of input-output pairs *k*_*max*_. Data from each individual’s four fMRI scan sequences are constructed in this manner, forming a dataset for each individual.

To illustrate the model’s prediction of whole-brain states, we used a three-node system as an example (Figure 6). For each pair of nodes (brain regions), the model takes as input the 10-TR fMRI time series from both regions and their structural connectivity, computing the interaction information via *I*(*x*). Using this interaction information and the original node states, the predicted state for the next time point in each region is computed with *R*(*x*). The predicted states are then compared to the actual fMRI values for the subsequent 10 TRs using the Huber loss, which is backpropagated to update the model parameters.

**Figure 6.**
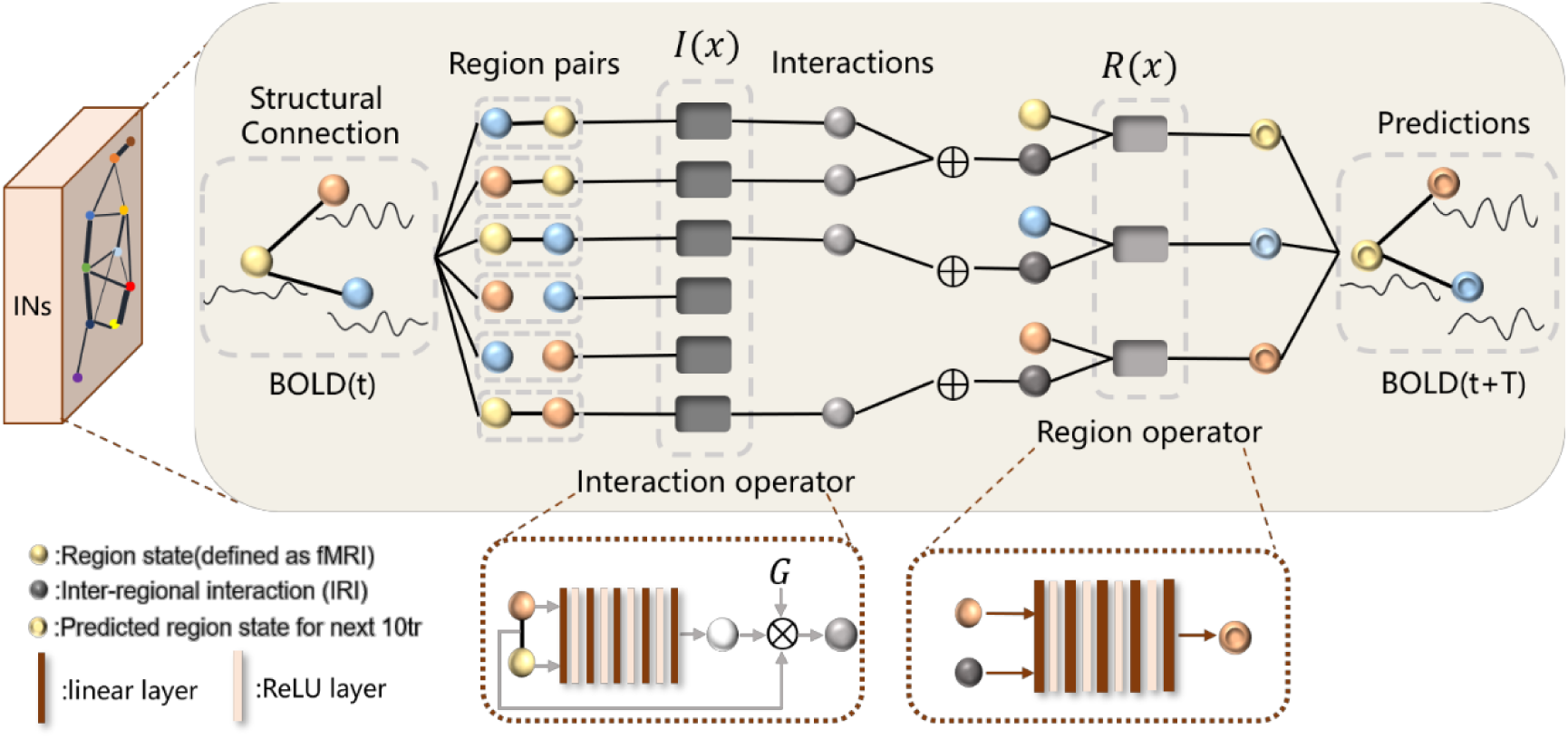
The overall operational process of the model. The information produced by interactions between brain regions is estimated using the interaction operator *I*(*x*), and the predicted values of the node state variables are estimated using the region operator *R*(*x*). Taking a three-node network as an example, for the red, yellow, and blue ball representing the brain region nodes, each pair of connected nodes inputs into the interaction operator to obtain the inter-region information (grey ball, *IRI*_*i*,*jj*_), a 30-dimensional vector. For each node, the total sum of IRIs on all the edges directly connected to it is computed as the total input information for that node (black ball). This total input information, along with the node’s own signal value, is entered into the region operator, yielding the predicted values of the brain region’s dynamic signals for the next 10 time points (concave ball). In the interaction operator, the red and yellow balls represent the signal values of two different brain regions with a connection, and the white ball is the output value of the multilayer perceptron. By multiplying this output with the connection weights (the lines between the red and yellow spheres) and the global parameter, the information value for the interaction between the two brain regions is obtained (grey sphere). In the region operator, the fMRI signal from the brain region itself, along with the information input the region receives, is fed into a multilayer perceptron to compute the region’s subsequent signals.

The dataset is split into training, validation, and test sets in an 8:1:1 ratio, with the initial 90% of the data serving as the training and validation sets and the final 10% designated as the test set. During training, the data for both the training and validation sets are randomly shuffled and re-partitioned at each epoch. We use torch_geometric to realize the model and load train data and the optimizer employed is Adam. The loss function utilizes Huber Loss(103) incorporating L1 regularization, which is a loss function that performs well across a variety of fitting tasks(104). The weight of regularization is 1e-4.The dataset is split into training, validation, and test sets in an 8:1:1 ratio, with the initial 90% of the data serving as the training and validation sets and the final 10% designated as the test set. During training, the data for both the training and validation sets are randomly shuffled and re-partitioned at each epoch. We use torch_geometric to realize the model and load train data and the optimizer employed is Adam. The loss function utilizes Huber Loss(103) incorporating L1 regularization, which is a loss function that performs well across a variety of fitting tasks(104). The weight of regularization is 1e-4.

Before training, we first randomly select a single individual for model hyperparameter optimization, and this individual does not participate in subsequent training and analysis. During the hyperparameter tuning process, global parameters are set based on previous research experience, with a value of *G* = 0.6. Grid search is used to optimize model parameters, including the dimension of information transferred between two brain regions, the output dimension of *I*(*x*), *Dim*_*IRI*_ ∈ [20,30,50,200], the number of layers in *I*(*x*), *L*_*I*_ ∈ [2,3,4,5,6], the number of layers in *R*(*x*), *L*_*R*_ ∈ [2,3,4,5,6], learning rate *Lr* ∈ [1 × 10^−3^, 5 × 10^−4^, 1 × 10^−4^, 5 × 10^−5^], weight decay *Wd* ∈ [1 × 10^−6^, 1 × 10^−7^, 1 × 10^−8^], and total training epochs *epochs* ∈ [50,100,300,400,1000,2000]. The model is selected based on the mean absolute error on the test set, with the final hyperparameters set as *Dim*_*IRI*_ = 30, *L*_*I*_ = 5, *L*_*R*_ = 5, *Lr* = 1 × 10^−4^, *Wd* = 1 × 10^−8^. Taking both training effectiveness and time into consideration, the number of epochs is set to 100.

### Model Validation

#### Mean absolute error

The mean absolute error (MAE) serves as a direct measure of the dissimilarity between predicted values and true values. Let the predicted value at time *T* be denoted as X_*T*_, and the true value as *Y*_*T*_, then X_*T*_ = [*x*_*T*,1_, *x*_*T*,2_, …, *x*_*T*,*N*_] and *Y*_*T*_ = [*y*_*T*,1_, *y*_*T*,2_, …, *y*_*T*,*N*_], where *N* = 360 is the total number of brain regions, *x*_*T*,*i*_ represents the predicted fMRI signal value for brain region *i* at time *T* and *y*_*T*,*i*_ represents the true fMRI signal value for brain region *i* at time *T*. The MAE is calculated by computing the absolute error across different brain regions and then taking the average of these absolute errors. The specific calculation process is defined by the following equation:

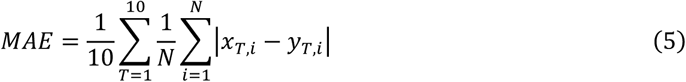

The above expression represents the MAE for a single prediction result. To ensure more reliable outcomes, we repeated the process 20 times on the test set, carrying out a total of 200 time points of fMRI signal prediction tasks, and calculated the average MAE to evaluate the model’s predictive performance, as showed in equation (6).

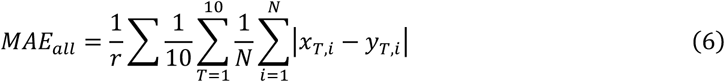

In the above equation, *r* = 20 denotes the number of times the computation is repeated. To compare the differences in the fitting effects across various brain regions, this study calculated the average MAE for different brain regions, as shown in equation (7).

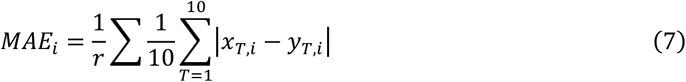

In equation (7), *MAE*_*i*_ represents the average MAE for the i-th brain region.

#### Continuous /segmented function connectivity

In this study, functional connectivity (FC) is obtained by calculating the Pearson Correlation Coefficient, whose formula is as follows:

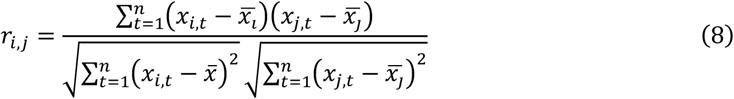

The term *r*_*i*,*jj*_ represents the correlation between the fMRI signal generated by brain region *i* and that by brain region *j*, where *x*_*i*,*t*_ denotes the fMRI signal value produced by brain region *i* at time point *t*, and *x*_*j*,*t*_ denotes the fMRI signal value produced by brain region *j* at time point *t*. The variable n represents the total number of time points used for the calculation. By performing the aforementioned operation between each pair of defined brain regions, the functional connectivity matrix between the brain regions can be obtained.

To obtain FC, this study employed two methods for generating the predicted sequences: continuous prediction and segmented prediction. Continuous prediction refers to inputting a 10 TR (time resolution) window, using the output as the next input to obtain the prediction result for the following time window, and then connecting these predictions to form a complete fMRI sequence. Let *χ*_*i*_ represent the input sequence, 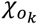 denote the output obtained from the k-th iteration, and *Se*_*pred*_ signify the output sequence; this process can be expressed by equation (9):

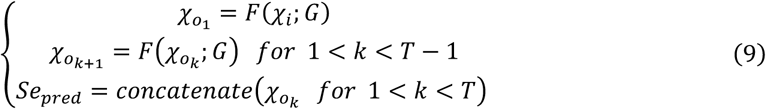

In equation (9), *F*(*x*) is the predictive model, *G* is the brain network, *T* is the number of cycles set, which is 10 in this study, and concatenate is a function that joins sequences. For functional connectivity validation, *χ*_*i*_ is set to the first 10 TR time window from the test set.

Segmented prediction refers to dividing a continuous time series into small intervals of 10 TR time windows, inputting each interval into the model to obtain the corresponding prediction, and connecting the different interval predictions chronologically to form a complete fMRI predicted sequence. Let the k-th interval be represented as 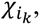 then 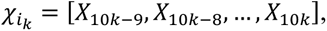 where X_*i*_ denotes the fMRI signal values for all brain regions corresponding to the i-th TR of the continuous time series. The prediction process is specifically shown in equation (10):

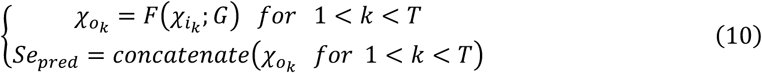

For this research, the chosen input is the continuous sequence before splitting the series that eliminates time points with head motion greater than 0.25. Starting from the 90th percentile position, a total of 100 TRs of the fMRI sequence is selected, and it is split into 10 groups with every 10 TRs as one group. The full sequence is thereby divided into 10 intervals, T=10.

The predicted sequences obtained from the two methods mentioned above both utilize the Pearson correlation method as outlined in equation (8) to calculate FC, which is then compared to the actual functional connectivity matrix. The actual sequence used for calculating the true FC is the one corresponding to the prediction input values; that is, in continuous prediction, the actual sequence starts from the 10 TR fMRI signal used as input and continues for a total of 100 TRs of fMRI sequence values, whereas in segmented prediction), the actual sequence is composed of the actual values corresponding to each segment’s output values. After obtaining the true FC and the predicted FC, the Pearson correlation coefficient is used to determine their correlation. The predicted FC is defined as *FC*^*pred*^ and the actual FC as *FC*^*truth*^. For *k* ∈ [*pred*, *truth*], 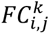 denotes the element in the i-th row and j-th column of the FC matrix, representing the functional connectivity of the fMRI signal between the i-th and j-th brain regions. The PCCs (Pearson correlation coefficients) between the predicted and actual FC can then be determined by equation (11).

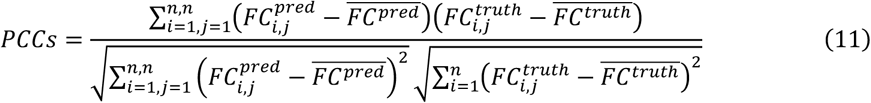

The equation (11) specifies that 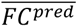 and 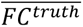 represent the global average values of the predicted and actual FC matrices, respectively.

### Model Analysis

#### Inter-regional interaction information (IRI)

Following model training, initial input data for the testing set were used to generate the model outputs. As described in the Methods-Models section, input data segmented every 10 tr yield a 30-dimensional message. By employing overlapping segmentation, a total of 200 such segments (each comprising 10 tr) were inputted, resulting in message data with dimensions of 30 × 200 × *N*_*sub*,*edge*_, where *N*_*sub*,*edge*_ indicates the total number of connectivity edges across subjects. PCA was applied to reduce the dimensionality of the obtained data’s first component, producing IRI data with dimensions of *N*_*pc*_ × 200 × *N*_*sub*,*edge*_, where *N*_*cp*_ is the number of selected components. In this experiment, most subjects’ first principal component (PC1) explained nearly 90% of the variance, so all IRI analyses were based on the dimensionality reduction to PC1, which means *N*_*pc*_ = 1.

#### IRI sent by region

For the IRI sent by region (RS-IRI), we first summed, for each region, the message data across all edges connected to that region along the corresponding dimensions. We then performed PCA on these region-level message vectors to derive the RS-IRI with dimensions of *N*_*pc*_ × 200 × *N*_*region*_. We used the MMP360 atlas so *N*_*region*_ = 360. The PC1 accounted for nearly 90% of the total variance, so we adopted PC1 as the RS-IRI, which means *N*_*pc*_ = 1.

### Model Application

#### Fast Fourier Transform (FFT)

To characterize the oscillatory patterns underlying dynamic inter-regional brain interactions, we applied Fast Fourier Transform (FFT) to the inter-regional information measures to obtain their spectral representations. Specifically, for each individual and each structural connection, the temporal inter-regional interaction measure *IRI*(*t*) was subjected to the Fourier transform as specified in Equation (12), yielding the frequency domain distribution *IRI*(*k*):

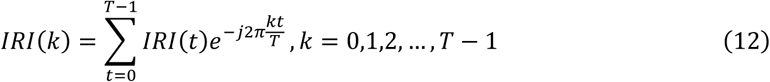

where *T* represents the total number of time points. In this study, the analysis was performed on *IRI*(*t*) data aggregated over 100 TRs, thus *T* = 100. Due to the inherently low sampling rate of the data, spectral analysis was confined to the frequency band of 0.005–0.5 Hz.

#### Group-level comparison between AD patients and controls

For all subjects, we first identified and extracted the common inter-region brain connections present across the entire cohort. The FFT was applied to the time series of these connections for each individual. For each edge (connection) and for each frequency band, the amplitude values were subjected to a two-sample t-test to compare the AD and NC group. The resulting p-values from these tests were then corrected using the FDR. Finally, we counted the number of inter-regional brain connections that still exhibited a significant group difference in each frequency band. These connections represent the pathways where the information transfer between the corresponding brain regions shows a significant group-wise difference between the AD and NC groups at that specific frequency band.

#### The association between oscillation amplitude and MMSE scores

First, inter-regional brain connections common to all subjects were identified. For each patient in the AD group, the FFT was applied to the time series of these connections. We then calculated Pearson correlation coefficients between the spectral amplitude of each edge within each frequency band and the participants’ MMSE scores. Statistical significance was assessed using a permutation test with 1,000 iterations. To control for multiple comparisons, p-values were adjusted using the FDR. Only results retaining significance after FDR correction are visualized in the figure.

## Author contributions

Conceptualization: SYL, DBZ, SXL

Methodology: SXL, DBZ, TTC

Investigation: SXL, DBZ

Visualization: SXL

Supervision: SYL

Writing—original draft: SXL, JCZ, ZKY, JJ

Writing—review & editing: SYL, SXL, DBZ, XXD, YRH, LC

## Data Availability Statement

The data used in this study are available through the Human Connectome Project (HCP; https://www.humanconnectome.org/study/hcp-young-adult/document/extensively-processed-fmri-data-documentation), a publicly available repository. The code and other data used for the analyses will be made publicly available on GitHub upon acceptance of the manuscript.

